# Proton-gating of opioid receptors and isoform functionality

**DOI:** 10.64898/2026.05.26.727816

**Authors:** Kyutae D. Lee, Gorav Surana, Sam Taylor, Daniel G. Isom

## Abstract

The three opioid receptors, mu (mu-OR), delta (delta-OR), and kappa (kappa-OR), are key G-protein-coupled receptors (GPCRs) involved in mediating pain relief. Yet, there has been a lack of understanding regarding how changes in the extracellular pH associated with inflammation affect their signal transduction pathways. This study demonstrates that all opioid receptors act as proton-gated coincidence detectors. A humanized yeast-based platform called DCyFIR, which isolates the pH environment outside the cell from potential confounding factors inside the cell, was employed to examine the activation of opioid receptors over a range of physiologically relevant to pathologically low pH environments. As such, the results show that agonist efficacy decreases with decreasing pH. This indicates that the receptor itself contains one or more protonatable sidechains that act as an intrinsic “pH sensor”. To determine the structural basis of this pH-sensing mechanism, structure-based pHinder calculations were used to identify potential pH sensing residues. Subsequent variant profiling identified the protonation of one conserved aspartic acid residue, D149 (also known as D3.32), located in a region near the binding site of opioid ligands, which, when exposed to acidic pH conditions, resulted in the disruption of the salt bridge necessary for effective docking of opioid ligands with the receptor. Consequently, the receptor lost all ability to be activated by agonists. Further characterization using a variety of functional assays on more than twenty different mu-OR alternative splicing variants indicated that all isoforms that possessed a continuous 7-transmembrane domain retained this proton-sensing capability. These studies collectively provide insight into the structural mechanisms underlying pH-dependent opioid receptor activation and lay the foundation for the design of new generations of selective analgesics that activate opioid receptors only in acidic inflammatory tissue.

## Introduction

Despite their potential for addiction and harmful side effects, opioids currently provide the best therapeutic option for treating chronic pain. Many efforts have focused on designing opioid drugs that circumvent adverse effects. Still, any major advances have been hindered by our limited understanding of opioid receptor signaling at the molecular level and by the translatability of findings between human and mouse models^1,2^. Opioids are agonists for the mu-, delta-, and kappa-opioid G protein-coupling receptors (GPCRs), with the mu-opioid receptor (mu-OR) being the primary target for pain relief, anesthesia, and suppressing diarrhea^3^. Although much is known about MOR signaling, many significant questions and challenges remain.

Recently, we have shown that many GPCRs are regulated by coincident acid signals (Fig. 1a^4^. These findings have ramifications for GPCR function across many biological contexts, such as acidified endosomes and the inflammatory zone^5^. Our study shows that the entire opioid receptor family is also regulated by pH. This discovery, as well as previous work showing that pH can regulate mu-OR agonist charge and binding^6–11^, holds great potential for the rational design of pH-intelligent therapeutics that target mu-OR in acidotic scenarios and exploit endosomal pH to control mu-OR signaling, recycling, and tolerance^5^. To succeed in this endeavor, we must first identify the molecular mechanism(s) of mu-OR pH sensing, including the residues that mediate mu-OR pH responses and the effects of pH on the charge state(s) of mu-OR agonists and inhibitors.

**Figure 1.**
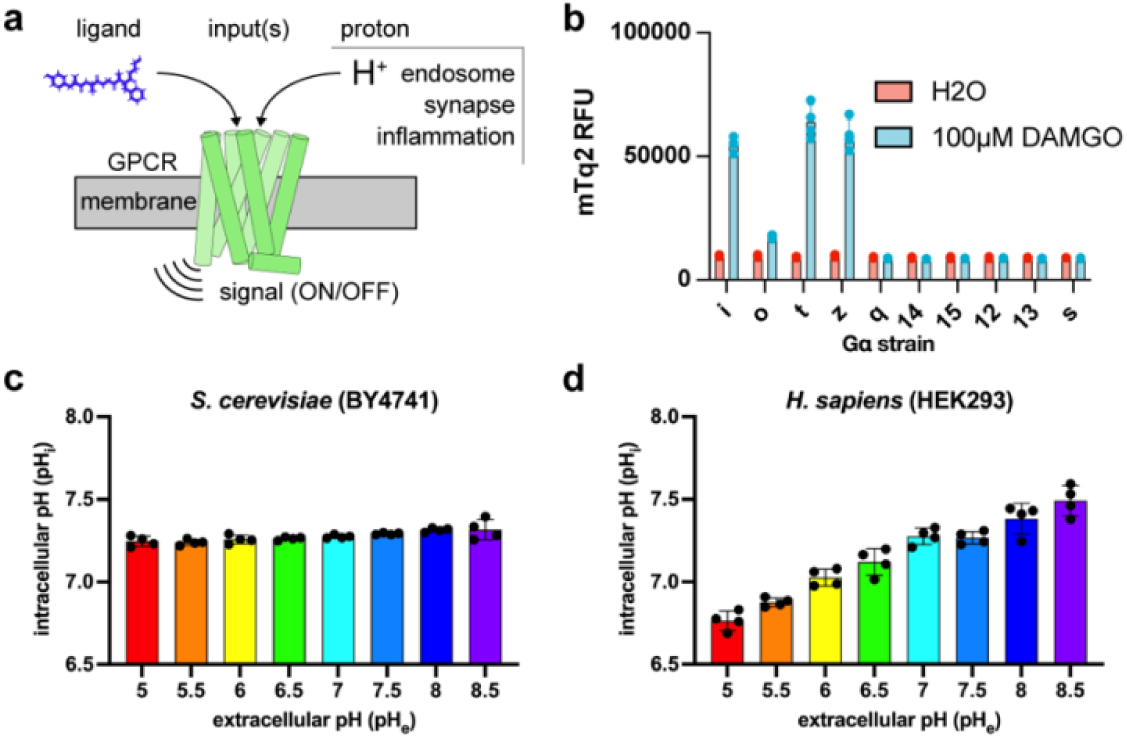
The DCyFIR humanized yeast system can isolate the receptor-level proton-sensing signal from intracellular pH-dependent confounders. (a) schematic representation of how GPCRs combine orthogonal ligand and extracellular proton inputs to make signaling decisions. (b) all ten Gα coupling profiles of the human mu-opioid receptor (mu-OR) in the DCyFIR humanized yeast system. mTq2 relative fluorescence units (RFU) at pHₑ = 7.0 stimulated with vehicle (H2O) or 100μM DAMGO. (c) intracellular pH (pH_i_) of BY4741 *S. cerevisiae* wild-type strain assessed using the ratiometric pHluorin biosensor across an extracellular (pHₑ) gradient from 5.0 to 8.5. (d) pH_i_ of HEK293 cells assessed under the same gradient; pH_i_ tracks pH_e_ approximately linearly across the full range. Data represent mean ± SD of four independent biological replicates.

Although the importance of pH has been established in previous studies, a fundamental mechanistic question remains unanswered: it is unclear whether the receptor itself, the ligand itself, or both in concert act as the pH sensor. Both synthetic and endogenously derived opioid ligands, as well as the extracellular and transmembrane regions of opioid receptors, commonly contain amine groups with pKa values within the physiological range^12,13^. A major constraint on investigating the extent to which each contributes to the overall pH-dependent regulatory mechanism arises from traditional approaches using *in vitro* systems. In all human cell lines studied extensively, e.g., HEK293T, intracellular pH (pH_i_) readily equilibrates with extracellular pH (pH_e_) (ref). As such, when attempting to manipulate pH_e_ to evaluate the ability of receptors to gate in response to changing pH_e_, one is simultaneously affecting pH_i_. Furthermore, Gα subunits possess pH-sensitive side-chain networks that inhibit downstream signaling when cytoplasmic pH decreases^14–16^. Therefore, signal suppression observed in mammalian models exposed to low pH cannot be definitively linked to proton-gated receptor activation at low pH.

To determine receptor-level functionality independent of pH_i_, we employed a new tool for evaluating GPCR pharmacology: DCyFIR (Dynamic Cyan Induction by Functional Integrated Receptors^17^). Unlike mammalian cell systems, yeast effectively buffers against changes in pH_i_. Regardless of changes in pH_e_, yeast maintains a constant pH_i_. We use this property to express human opioid receptors paired with humanized Gα chimeras and measure GPCR activity levels in response to changes in pH_e_, independent of pHi. Utilizing structural informatics (pHinder)^18^, we identified a set of amino acids within the transmembrane domain of mu-OR that potentially contribute to sensing changes in pH_e_. To further elucidate how opioid receptors respond to pH variations *in vivo*, we profiled these variants to examine how pH regulates opioid receptor signaling output as coincidence-detection mechanisms.

Most human GPCRs have few isoforms. The second half of the study addresses this long-standing problem, which arises from the fact that modern opioid biology focuses solely on the canonical mu-OR isoform (isoform 1). In fact, there are at least 20 known MOR isoforms in humans and > 34 in mice. Noncanonical MOR isoforms are classified as 7 transmembrane (7TM), 6TM, or 1TM splice variants^19–22^ that can far exceed the abundance of the canonical MOR isoform in specific brain regions important to behavior and addiction, such as the prefrontal cortex, thalamus, and hypothalamus^23,24^. Additionally, prolonged opioid use can cause the levels of MOR isoforms to change dramatically. For example, the mouse 7TM isoforms mMOR-1A and mMOR-1B increase by 83-fold in the striatum, reducing the canonical isoform to only 30% of the isoform pool^23,24^. However, the functional relevance of these and the other MOR isoforms remains unclear, as they have yet to be systematically assessed in any cell-based system. The resulting splice variants represent unexplored opportunities to study basic aspects of opioid receptor biology, and this study employs the same yeast-based DCyFIR platform to characterize them systematically.

## Results

### Establishing Receptor-Level Proton Coincidence Detection with the DCyFIR Platform

We first had to establish opioid receptors in the DCyFIR platform. In our previous study^4^, opioid receptors were not included, but using high-copy yeast plasmids to increase expression uncovered opioid receptor signaling. Utilizing this platform, we assessed wild-type mu-OR across all ten yeast-human Gα chimeras that cover all sixteen human Gα (Fig. 1b). Full agonist DAMGO stimulation at 100 μM at pH_e_ of 7.0 elicited robust, target-specific Gα signaling from Gαi, Gαo, Gαt, and Gαz chimeras, staying true to what is known about mu-OR Gα specificity.

To test whether opioid receptors encode intrinsic proton-sensing properties into their structure, we first had to eliminate the confounding variables associated with traditional mammalian expression platforms. HEK293T cells lack robust buffering mechanisms to maintain intracellular pH when exposed to external pH gradients. Therefore, exposing HEK293T cells to an extracellular pH gradient ranging from 7.5 to 5.0 resulted in an intracellular pH drop to ∼6.7 at pH_e_ 5.0 (Fig. 1c). To address these limitations, we utilized the DCyFIR platform, where yeast (S. cerevisiae) maintains cytosolic pH through highly efficient proton pumps, resulting in no change in intracellular pH even when exposed to an extracellular pH of 5.0 (Fig. 1d). By stabilizing the intracellular pH baseline, any GPCR output changes can be assigned to extracellular protonation events versus intracellular acidification.

### Acidic Extracellular Environment Dynamically Changes Clinical Opioid Pharmacologies

To assess how environmental acidification affects the pharmacological characteristics of opioids, the pKa of each opioid dictates its protonation state at physiological pH (7.4) and at an acidotic, inflamed microenvironment pH (6.5). The pKa of opioid agonists and antagonists is shown in the table in Fig. 2b. Employing the Henderson-Hasselbalch equation, the active, protonated fraction shifts from 1.2% to 9.1% for β,β-difluoro fentanyl (6b), from 33.4% to 79.9% for DAMGO, and from 86.3% to 98.0% for PZM21 (Fig. 2a,b). Therefore, an opioid like PZM21, morphine, and β,β-difluoro fentanyl (6b) would not significantly change its protonated state in each scenario, whereas opioids like DAMGO and the endogenous peptide agonists (enkephalins) would (Fig. 2a).

**Figure 2.**
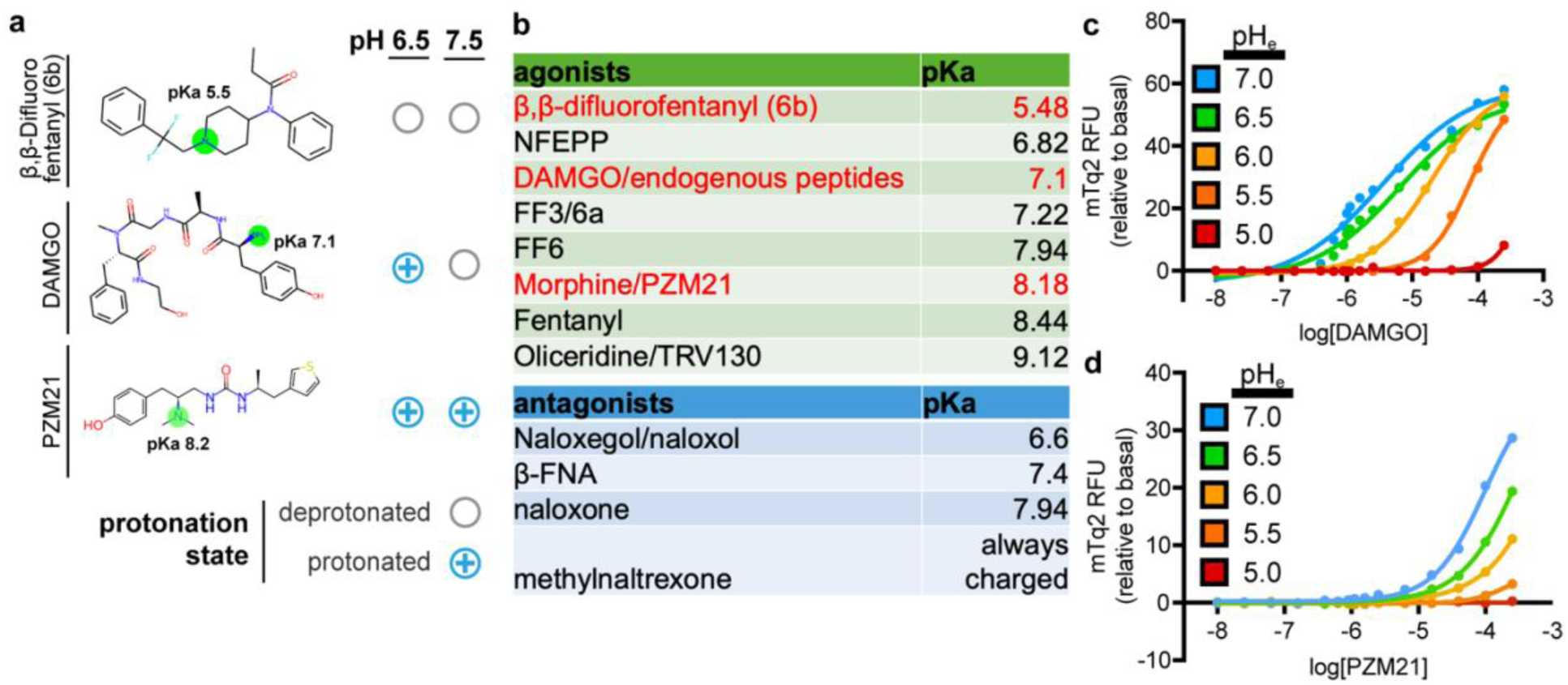
Protons modulate ligand pharmacology of mu-OR depending on the ionizable properties of ligands. (a) representative structures of three opioid ligands spanning the pKa range studied here, with the ionizable amine group of each ligand highlighted in green. The predominant protonation state of each ligand at pH 6.5 and pH 7.5 (filled circle, protonated; open circle, un-protonated). (b) tabulation of common and experimental opioid agonists (top, green) and antagonists (bottom, blue) with pKa values of the relevant ionizable amine group. Agonists highlighted in red are the representative ligands shown in (a). (c-d) dose response curves for (c) DAMGO and (d) PZM21 for mu-OR signaling in the DCyFIR humanized yeast system at pH_e_ levels of 5.0, 5.5, 6.0, 6.5, and 7.0 with mTq2 reporter as readout of signaling. The signaling levels are normalized relative to basal at lowest concentration of ligand tested. Data presented are mean ± SD of four independent biological replicates.

We then established the dose-response relationships of DAMGO and PZM21 across a pH_e_ gradient from 7.0 to 5.0 (Fig. 2c,d). At all tested pH valu_es_, DAMGO activated mu-OR and reached peak efficacy (Fig. 2c). However, as the extracellular environment became increasingly acidic, potency decreased significantly. The PZM21 agonist showed a different degree of decline as the extracellular environment became more acidic, suggesting that, in addition to receptor-level protonation states, ligand protonation state also contributes to the rate of decline. While assessing the effects of other opioids in the table (Fig. 2b) would provide valuable mechanistic insight, we could not include these specific derivatives due to institutional licensing limitations.

### Identification and Mutational Analysis of Buried Ionizable Sensor Network

Having confirmed that opioid receptors serve as coincidence detectors, we next attempted to define the structural features underlying this behavior. We postulated that buried ionizable residues within the TM bundle undergo context-dependent protonation, thereby modifying the receptor’s conformational equilibrium. We identified a candidate sensory network in mu-OR consisting of D2.50 (D116), D3.32 (D149), H6.52 (H299), and H7.46 (H321) using pHinder^25^ (Fig. 3b). We chose to focus our analysis on two of these four nodes: D3.32 (D149) and H6.52 (H299). We left the remaining two nodes: D2.50 (D116) and H7.46 (H321) for future studies.

**Figure 3.**
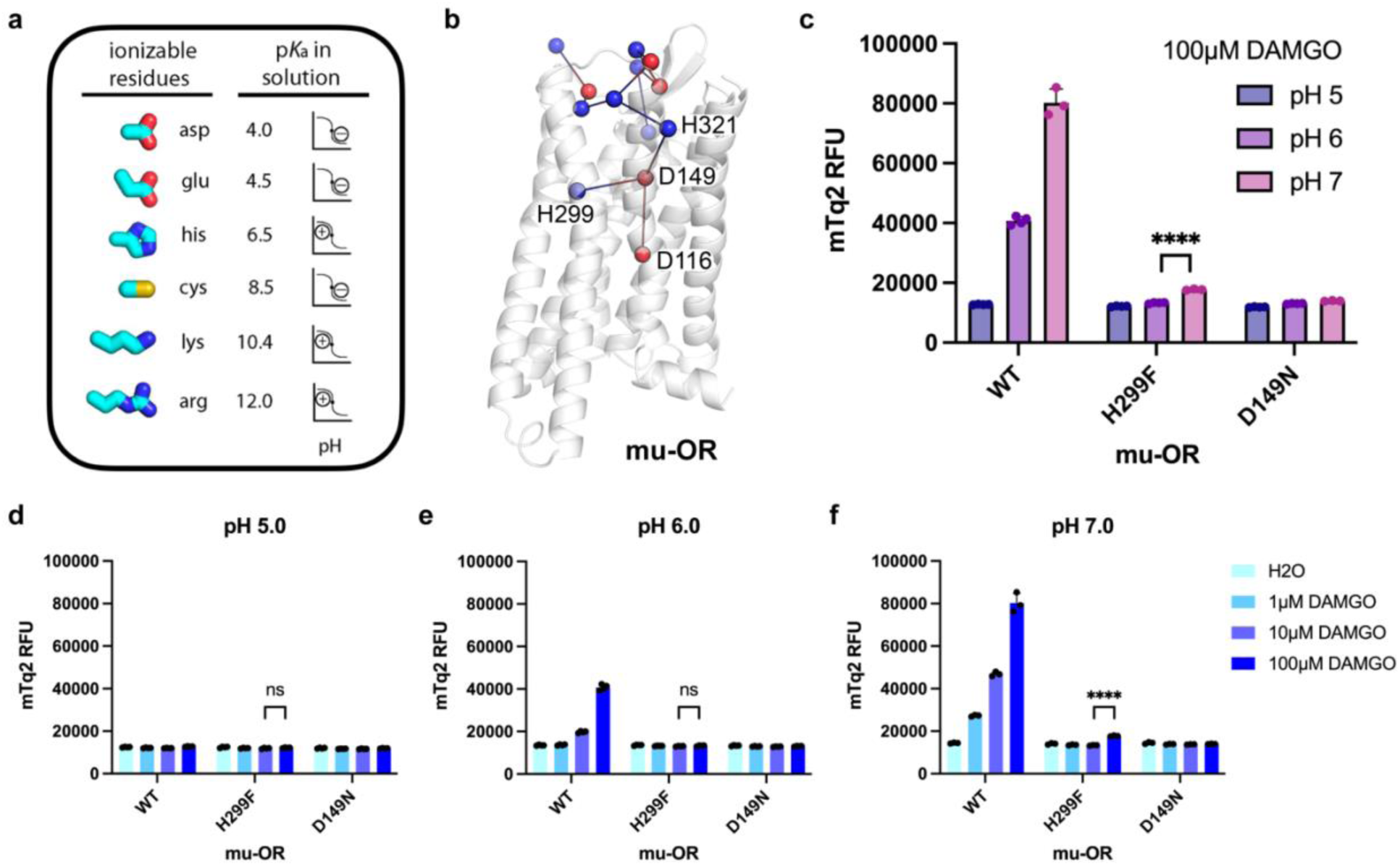
Agonist induced signaling is structurally encoded in mu-OR and requires a buried ionizable network. (a) reference panel for six potential ionizable side chains present in proteins along with their solution pKa values and titration behavior. (b) candidate residues identified by the pHinder algorithm in the mu-OR (PDB: 8EF5) transmembrane region for proton sensing. Acidic residues (D116, D149) are colored red and histidine residues (H299, H321) are colored blue. (c) Gαi-coupled DCyFIR signaling levels of wild-type (WT) mu-OR and the two point mutants H299F and D149N at various pH_e_ levels stimulated by 100μM DAMGO. (d-f) Gαi-coupled DCyFIR signaling levels for DAMGO for wild type, H299F and D149N mu-OR mutants at indicated concentrations of DAMGO in pH_e_ of 5.0, 6.0, and 7.0. Data presented are mean ± SD of four independent biological replicates (****, p<0.0001 t-test for specified comparisons; ns, not significant).

D3.32 is known to interact with the protonated amine moiety present on most opioid ligands, including morphinans and peptides, through an electrostatic salt bridge^26–28^. Although this interaction is clearly essential for the binding of opioid ligands to mu-OR and subsequent receptor activation, it has not been determined if this interaction is regulated by pH. Computational models suggest that D3.32 resides in a sufficiently hydrophobic region to elevate its pKa to ∼6.5 from its natural pKa (Fig. 3a). Thus, at physiological pH (∼7.0), D3.32 would likely remain deprotonated and negatively charged, thereby preserving the salt bridge. Under more acidic conditions (below pH_e_ ∼6.5), where inflamed or ischemic tissues often reside, D3.32 would become increasingly protonated and thus lose its negative charge, disrupting this electrostatic anchor point and promoting movement toward an inactive conformation regardless of ligand concentration (Fig. 3c-f).

To examine these predictions experimentally, we created isosteric versions of mu-OR containing mutations at D3.32: D3.32N (D149N), which mimics the protonated form of native D3.32; and H6.52F (H299F), which removes an additional ionizable node within the TM bundle. According to our prediction, both substitutions should result in signaling incompetent receptors: D3.32N, which represents an inactive state indistinguishable from acid-off; and H6.52F, which removes an important ionizable residue involved in receptor activation. Our experimental results supported these predictions: mu-OR D149N lost all responsiveness to all agonists tested, including DAMGO, Met-Enkephalin, Leu-Enkephalin, SNC80, and asimadoline at all concentrations and pH values tested (Fig. 5c). Similarly, H6.52F substitution yielded a signaling incompetent phenotype (Fig. 4c). These results demonstrate that both D3.32 and H6.52 are essential for activating mu-OR and support our model wherein D3.32 undergoes protonation in low-pH environments and leads to suppression of receptor activation.

**Figure 4.**
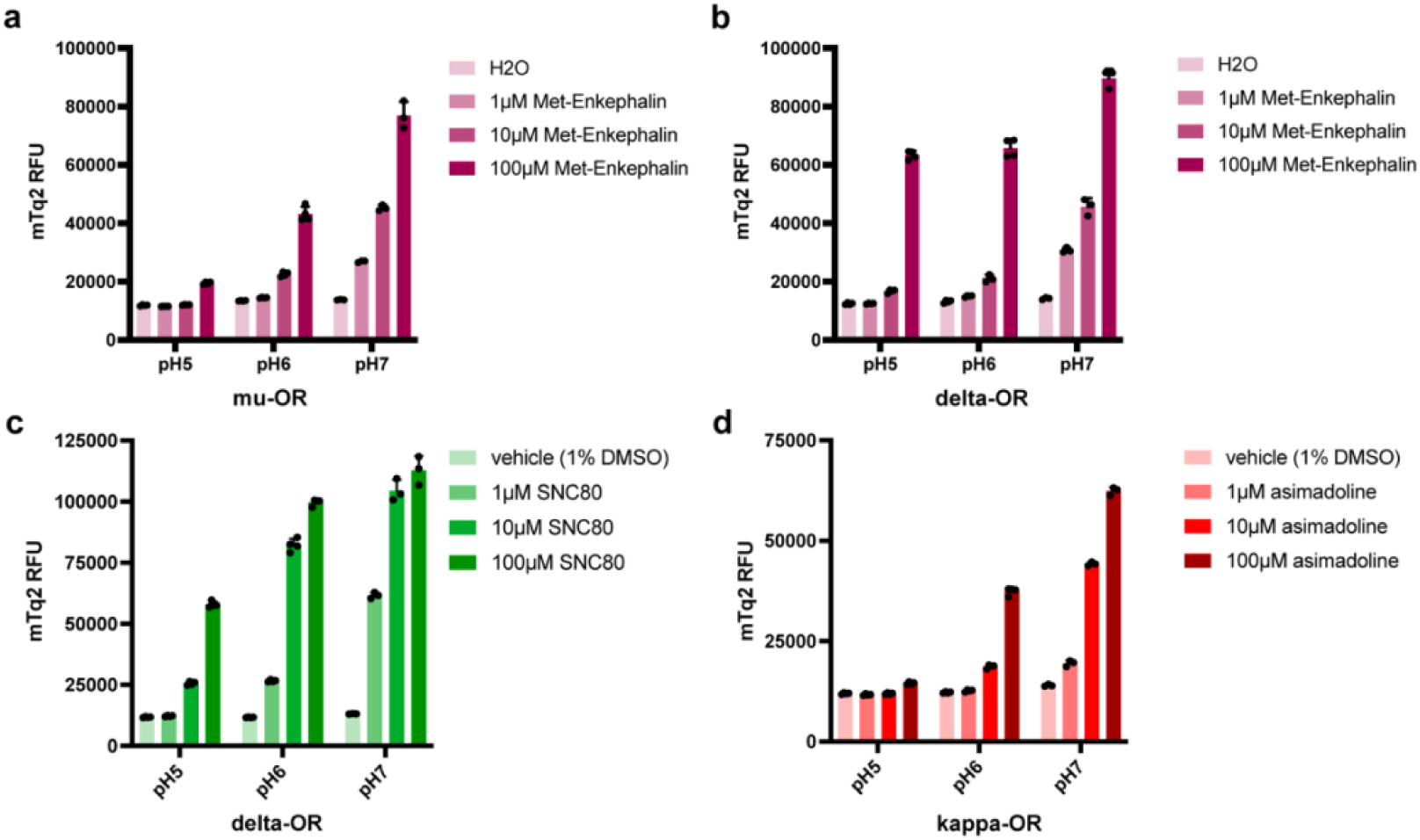
Receptors encode pH-gated opioid signaling that varies across subtype when stimulation is provided by the same ligand. (a-d) Gαi-coupled DCyFIR signaling levels in various pH_e_ of 5.0, 6.0, and 7.0 for opioid receptor and ligand pairs of: (a) mu-OR; met-Enkaphalin, (b) delta-OR; met-Enkaphalin, (c) delta-OR; SNC80, (d) kappa-OR; asimadoline. Data presented are mean ± SD of four independent biological replicates.

**Figure 5.**
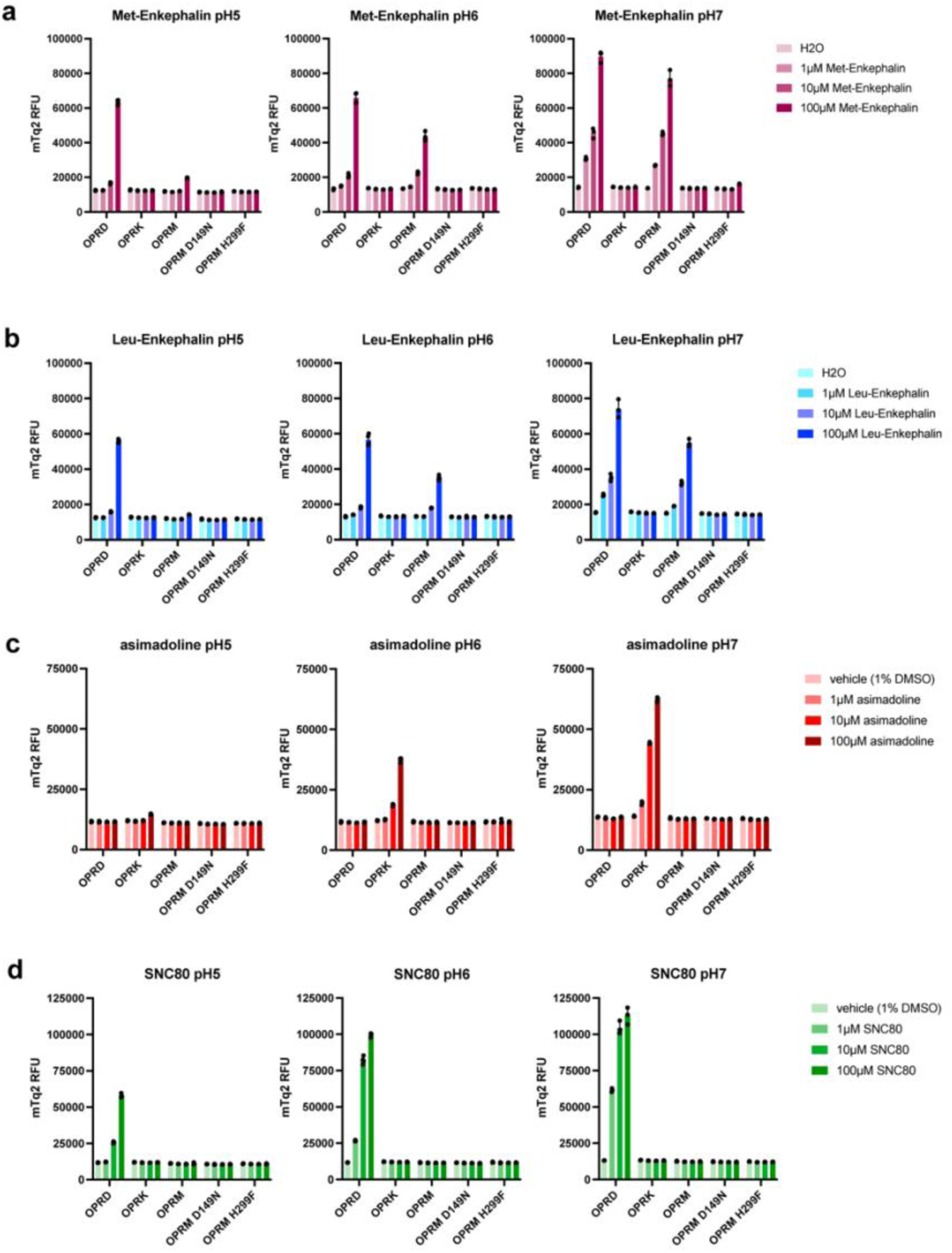
Cross-receptor and cross-mutant analysis reveals mechanisms encoding specific contributions to ph-gated opioid signaling. (a-d) Gαi-coupled DCyFIR signaling levels in various pH_e_ of 5.0, 6.0, and 7.0 for opioid receptor (wild-type OPRD=delta-OR; OPRK=kappa-OR; OPRM=mu-OR; and mu-OR variants D149N and H299F) and ligand pairs of: (a) met-Enkaphalin, (b) leu-Enkaphalin, (c) asimadoline, (d) SNC80. Data presented are mean ± SD of four independent biological replicates.

### Structural Requirements for Proton-Gated Behavior Differ Between Opioid Receptor Subtypes

To separate receptor-level pH-dependent behavior from ligand protonation-related behavior, we directly compared the proton-gated behaviors of mu-OR, delta-OR, and kappa-OR activated by the same concentrations of the same ligand. We selected Met-Enkephalin as our ligand because it is a natural peptide that selectively activates delta-OR and moderately activates mu-OR (Fig. 4a). Met-Enkephalin activated both receptors to produce similar levels of signaling at pH_e_ 7.0 (Fig. 4b). However, as the pH_e_ was reduced, the two receptors behaved differently (Fig. 4c). The suppression of mu-OR activity was consistent with what was observed with synthetic ligands. Delta-OR activity remained robust at pH_e_ 5.0, but there was evidence of reduced affinity as evidenced by a rightward shift in the apparent EC50 value. Because Met-Enkephalin has the same protonation state regardless of which receptor it binds to, this differential behavior demonstrates that proton sensitivity is a structural feature encoded in the receptor itself rather than in the ligand alone. Similarly, we observed receptor-specific divergence with other synthetic agonists as well: asimadoline-stimulated kappa-OR activity was reduced by approximately 75% at pH_e_ 5.0, whereas delta-OR activity stimulated by SNC80 retained its high levels of activity under the same acidic conditions (Fig. 4d-e).

### Characterization of Structural Requirements for Coincidence Detection via Splice Variants

OPRM1 encodes over 20 naturally occurring splice variants *in vivo*^22,23,29^. To investigate how variations in structural organization affect competence for coupling to G-proteins, we performed a comprehensive functional screening of twenty naturally occurring splice variants of mu-OR using the DCyFIR platform (Fig. 6a).

**Figure 6.**
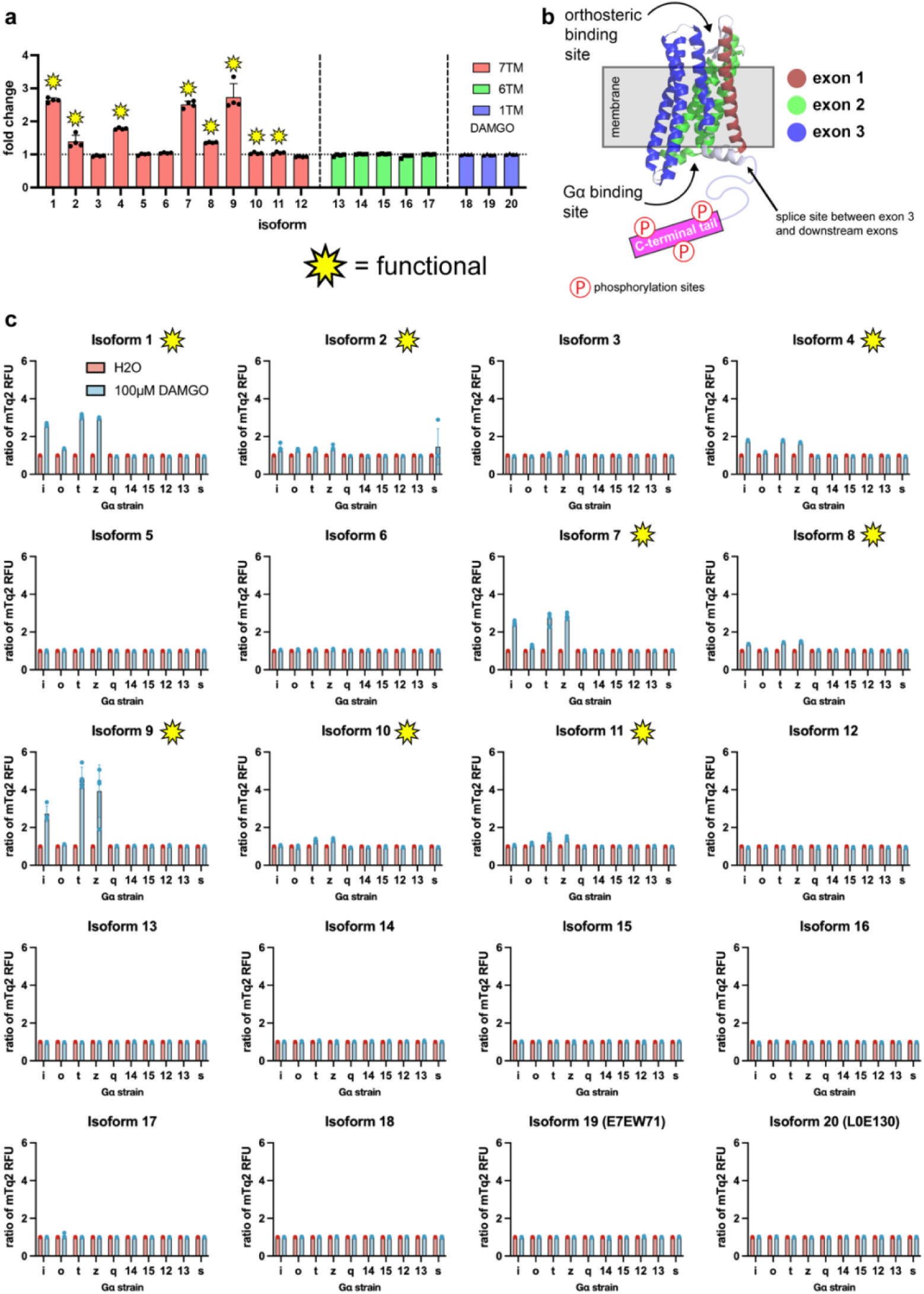
The signaling competency of splice variants for the OPRM varies and is confined to those having a full length 7-transmembrane domain. (a) A summary of agonist-induced signaling for twenty different human splice variant forms (isoforms 1-20) of mu-OR that were tested in the DCyFIR system. Bars indicate the magnitude of increase in mTq2 signal relative to vehicle after stimulation with 100µM DAMGO for each of the 20 isoforms tested. Variants are color coded according to their predicted topological characteristics: 7TM (red); 6TM (green); and 1TM (blue). Functional isoforms (isoforms 1, 2, 4, 7, 8, 9, 10, 11) are marked with a yellow star as defined by its response to DAMGO coupled to any Gα. The splice variants possessing either a reduced topology (6TM) or severe truncation (1TM) did not produce measurable responses to DAMGO treatment. (b) Schematic illustration of mu-OR topology showing the three exons that code for the receptor (exon 1: red; exon 2: green; exon 3: blue), the location of the orthosteric binding site, the G-protein alpha subunit interaction surface and the C-terminal tail with phosphorylation sites that are important for. The splice site located immediately downstream of exon 3 where most of the isoform diversity profiled here was generated through alternative splicing. (c) Each graph represents each of the isoform G-protein coupling of all twenty isoforms tested against each of the ten Gα in the DCyFIR system. Each bar indicates the ratio of mTq2 RFU measured upon incubation with 100µM DAMGO over vehicle (H2O). Data points represent mean ± SD values from n = 4 independent biological replicates.

Activation of these isoforms with DAMGO at a concentration of 100μM at pH_e_ 7.0 revealed that only eight isoforms that retain full-length seven-transmembrane (7TM) architectures demonstrated robust agonist-dependent Gα chimeric protein coupling: isoforms 1, 2, 4, 7, 8, 9, 10, and 11 (Fig. 6a,b). All remaining isoforms failed to exhibit agonist-dependent Gα coupling at any agonist concentration. Mapping nonfunctional isoforms onto the canonical GPCR topology indicated that all signaling competent isoforms preserve a full-length seven-transmembrane bundle (Fig. 7a,b). Isoforms lacking full-length seven-transmembrane bundles as a result of either C-terminal truncations, reduction in the number of TM domains, or extensive C-terminal deletions stemming from premature splice junctions between exon three and downstream exons did not engage in Gα-coupling.

**Figure 7.**
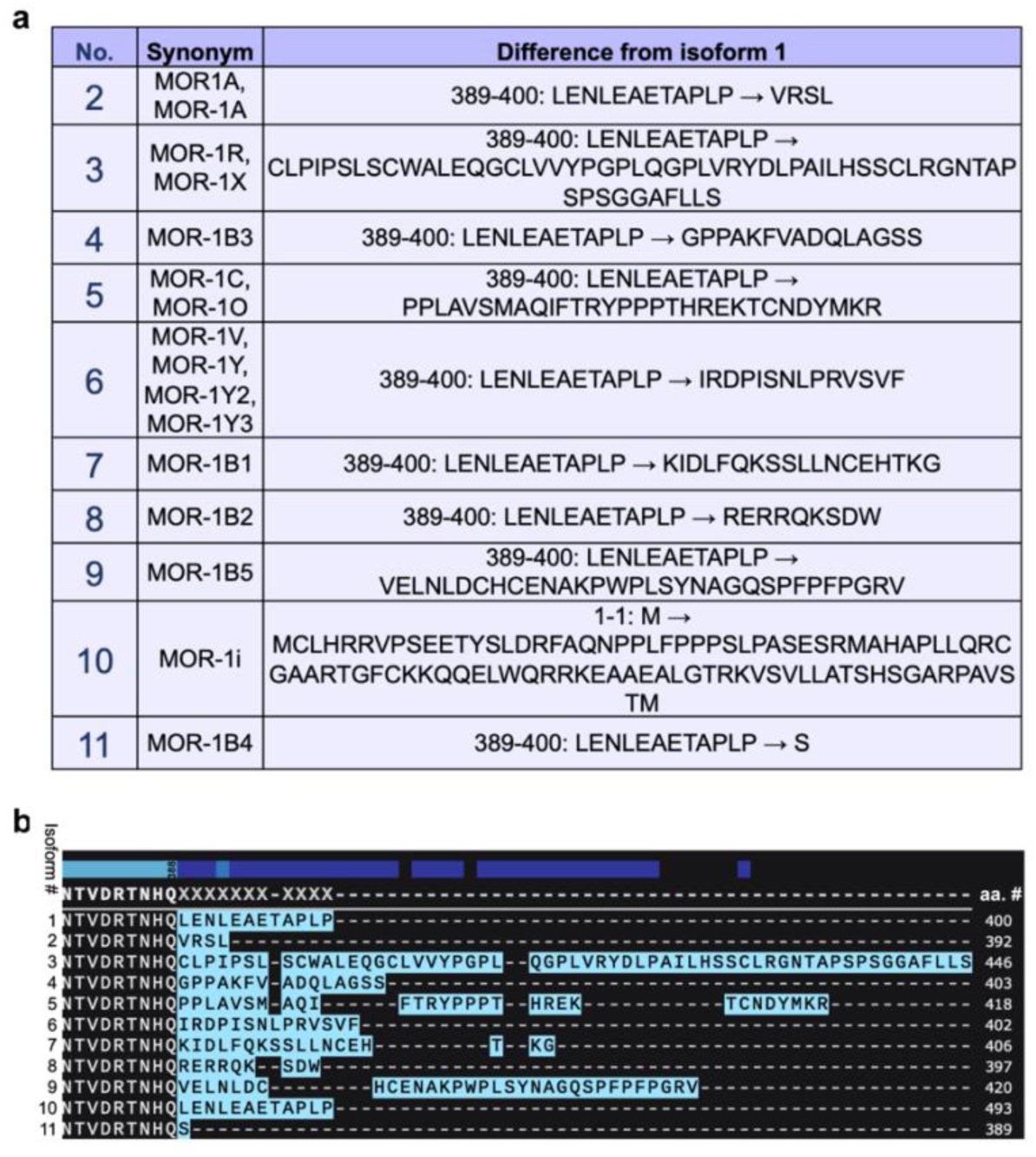
Sequence differences underlying all human OPRM splice variant diversity. (a) a summary table illustrating the ten other 7TM containing OPRM isoforms (isoforms 2-11), with isoform 1 being the reference full-length receptor. Numbers in the third column indicate the amino acid positions in isoform 1. Listed for each isoform are one or multiple receptor synonyms commonly employed in the literature (e.g. MOR-1A, MOR-1B series, MOR-1R, MOR-1C, MOR-1V/Y/Y2/Y3, MOR-1i) and the specific sequence difference of that isoform compared to isoform 1. Only isoform 10 (MOR-1i) differs in sequence from isoform 1 at the N-terminus, whereas all other isoforms vary in sequence at the C-terminal tail. (b) Alignment of sequences corresponding to the divergent C-terminal regions of isoforms 1-11. Highlighted at left is the upstream sequence “anchor” that is common to all five of these isoforms (NTVDRTNHQ-388) followed by unique residues present within each isoform. Dashes (-) indicate positions where gaps exist due to differences in length of each isoform. The last residue number assigned to each isoform is noted on the right side of the table and illustrates how significant a role alternative splicing plays in determining receptor length variability.

These results are consistent with previous pharmacological characterization studies conducted by the Pasternak group^29^, demonstrating that, for classical opioid analgesia, full-length seven-transmembrane receptors are required for canonical opioid receptor signaling pathways. While our results do not exclude alternative functions for noncanonical signaling-difficient isoforms, previous studies (MOR-1B2 and mMOR-1B4, implicated in different Gα specificities^30^) have suggested several examples of alternative roles for some of these isoforms as scaffolding proteins for heterodimers or as pharmacological “sinks”. The results here indicate that none can engage agonist-mediated activation in isolation.

## Discussion

The primary purpose of the research presented in this dissertation was to explore the structural and functional mechanisms underlying pH-dependent signaling in the opioid receptor family. Using three opioid receptors –– mu-opioid receptor (MOR), delta-opioid receptor (delta-OR), and kappa-opioid receptor (kappa-OR) –– we found that they serve as proton-gated coincidence detectors. Each opioid receptor’s output depends on the simultaneous chemical environment of the external microenvironment, not just on the presence of an agonist. Mapping these capabilities to a network of buried ionizable residues distributed throughout the transmembrane core provided the first description of how these receptors integrate their local environment into their signaling process.

However, before the structure-function relationship could be defined, a confounding issue had to be addressed. Conventional receptor pharmacology does not account for the fact that HEK293 cells’ internal pH tracks changes in their external pH, nor does it consider that G protein alpha subunits also act as pH sensors. Therefore, when using mammalian models to evaluate the response of opioid receptors to changes in external pH, the observed decrease in receptor activity results from a combination of receptor gating and G-protein-mediated inhibition that cannot be easily distinguished. The yeast-based DCyFIR system circumvented this problem. The near-neutral internal pH of *S. cerevisiae* under acidic conditions (Fig. 1a-b) made it possible to decouple the effects of protonation of the opioid receptor itself and subsequent G-protein inhibition.

Structural bioinformatics studies have previously identified D2.50(D116), D3.32(D149), H6.52(H299), and H7.46(H321) as potential candidates for proton-sensing networks in mu-OR. Functional studies have evaluated two of these sites. It has been well documented that D3.32 (D149) plays a key role in facilitating the electrostatic interactions required for opioid ligand binding^26–28^. What has not previously been reported is that this interaction is pH-dependent. Based on prior studies, we propose that the hydrophobic nature of D3.32 increases its pKa to approximately 6.5^31,32^. Under acidic conditions, progressive protonation neutralizes the anionic charge associated with D3.32, disrupts the salt bridge formed with the amine portion of the bound ligand, and results in a shift towards the inactive conformation of the receptor. Substitution of D3.32N resulted in a loss of function across all tested ligands and pH values (Fig. 4c); based on the proposed mechanism, we predict that this would occur regardless of whether the receptor was activated or inhibited. These results confirm that receptor activation is conditional on the ionized state of D3.32. Similar results were obtained using the H6.52F substitution. Functionally testing D2.50 (D116) and H7.46 (H321) would be informative in future studies.

In addition to defining a common mechanism for proton sensing among various GPCRs, this study also has a related parallel with an accompanying study examining A2AR (bioRxiv). In that work, D2.50 and N7.49 were identified as the functional units responsible for A2AR-mediated proton-sensing, and substitution of these residues with either isosteric amino acids or alanines produced pH-insensitive rather than signaling-deficient variants (i.e., mutations that did not abolish receptor signaling but rather abolished pH dependence). There is significant interpretative value in making this distinction. For example, although mutation of D2.50(D116) did not occur in this study, mutation of D3.32(D149) eliminated receptor function. Whether these differences reflect fundamentally different structures supporting similar mechanisms or simply different network positions (e.g., D3.32 in mu-OR vs. D2.50 in A2AR) is currently unknown and will require detailed comparisons of structure and function to clarify.

Comparative receptor profiling provides the most compelling evidence that proton-sensitivity is inherent in the structural features of each receptor. As shown in Fig. 3(b-c), mu-OR and delta-OR exhibit disparate responses to identical Met-Enkephalin concentrations at pHe = 5.0: whereas delta-OR exhibits nearly maximal efficacy at this pH, mu-OR exhibits substantially reduced efficacy. Given that the protonation states of the ligand are independent of the type of receptor it binds to, this difference illustrates that each receptor type possesses a unique structural propensity for proton-sensitivity. While we have defined the structural elements of pH-sensitivity in mu-OR, we have not done so in delta-OR or kappa-OR. An examination of a sequence alignment of the four candidate proton-sensing positions across mu-OR, delta-OR, and kappa-OR reveals numerous non-conservative residues (Fig. S2). If a position essential for pH-gating in mu-OR (e.g., H6.52) is replaced with a non-ionizable residue in delta-OR, then this would provide a structural rationale for why delta-OR retains signaling competency at acidic pH conditions. Reciprocal substitutions in mu-OR and delta-OR, designed to test this hypothesis, are thus an immediate next step.

The pH-gated logic of opioid-receptor-mediated signaling described herein has implications for the design of context-specific analgesics. One of the major limitations of traditional opioids is their ability to activate mu-OR-containing neurons in both acidic (e.g., peripheral nerves) and neutral (e.g., brainstem/CNS) tissue environments. By exploiting this pH disparity at the level of ligand design, it may be possible to create drugs that selectively bind to and activate opioid receptors only at acidic pH sites (e.g., nociceptors) ^6,33^. Achieving this would require ligands whose own protonation state shifts their receptor affinity in the opposite direction. For example, NFEPP-style analogs whose protonated (binding-competent) form predominates only at low pH, partially compensating for the receptor’s intrinsic acid-off behavior. Furthermore, our structural identification of the D3.32 proton-sensing mechanism provides a complementary receptor-level paradigm. By engineering ligands whose binding affinities or efficacies depend on the protonation state of D3.32, it may be possible to develop drugs that exploit pH-dependent conformational changes in the transmembrane domain for selective binding.

The splice variant screen illustrated additional constraints on the spatial relationships required for coincidence detection. Only eight out of twenty natural mu-OR isoforms examined herein retain their full-length 7TM topology (isoforms 1, 2, 4, 7, 8, 9, 10, 11) and exhibited signaling competency in the DCyFIR assay (Fig. 5a-b). One limitation is that the expression level of these isoforms was not measured. With that in mind, these findings support pharmacological studies by the Pasternak group on canonical mu-OR isoform pharmacology and extend them by creating a direct functional map that illustrates which isoforms activate the Gα-dependent pathway used for coincidence detection. Additionally, these results indicate that the C-terminal tail clearly has demonstrable effects in G alpha coupling, not only for potential effects in arrestin recruitment. Establishing whether such selective binding affinities/efficacies can be achieved with sufficient practicality remains an unresolved question, and this structural and functional analysis will help address it.

## Methods

### Mammalian Cell Culture and Transfection

Human cell lines were obtained from ATCC and cultured under recommended conditions. HEK293T cells were cultured in DMEM (Thermo Fisher Scientific, #11995065) with 10% FBS (HyClone, #SH30071.03) and 1% Penicillin-Streptomycin-Glutamine (PSG, #10378016). All cells were incubated at 37 °C with 5% CO₂ and passaged using TrypLE Express (ThermoFisher, #12604013). For transient expression experiments, cells were seeded in 6-well plates and transfected using TransIT (Mirus Bio, #MIR 2305). After 24 h, cells were trypsinized, counted, and replated into white 384-well plates (Greiner Bio-One #781095) at 5,000–10,000 cells per well for downstream assays.

### Yeast DCyFIR Strains

Yeast strains used in this study are identical to those previously described^17^. Briefly, we genetically engineered the wild-type yeast strain BY4741 to include 1) deletion of the native yeast GPCR (ste2Δ) and its negative regulators of the pheromone pathway (GTPase activating protein sst2Δ, cell cycle arrest factor far1Δ); 2) installation of a CRISPR addressable expression cassette in chromosome locus X2 that includes a constitutive promoter (PTEF), synthetic universal targeting sequence for CRISPR/Cas9 editing (i.e., knock-in of human GPCRs), and terminator (TCYC1B); and 3) installation of the mTurquoise2 transcriptional reporter in place of the pheromone responsive Fig1 gene (fig1Δ:: mTurquoise2). After genetic modification of this base strain to remove these elements, we expanded the number of GPCR-Gα strains to ten through CRISPR-Cas9 genome engineering to add human C-terminal sequences to the native yeast Gα subunit Gpa1. Finally, we expressed human GPCR genes into each of the ten GPCR-Gα strains using the yeast 2μ high copy plasmid (pYEplac181). Additional information regarding the development of these yeast strains can be found in references^4,17^.

### Media

All yeast strains were streaked on YPD media plates (20g/L peptone, 10g/L yeast extract, 2% glucose, 15g/L agar) from a frozen 30% glycerol stock at –80°C. All DCyFIR screen experiments and pH titration studies utilized yeast that were cultured in low-fluorescence synthetic complete drop-out media (SCD media; 50 mM potassium phosphate dibasic, 50 mM MES hydrate, 5 g/L ammonium sulfate, 1.7 g/L yeast nitrogen base w/o amino acids, folic acid, and riboflavin (Formedium; CYN6505), 0.79 g/L complete amino acid mix (MP Biomedicals; 4500022), and 2% glucose) titrated to the desired pH by adjusting with HCl or NaOH. In addition to the use of SCD media for yeast growth in pHluorin-based studies where intracellular pH of yeast was assessed through measurement of pHluorin fluorescence, yeast were also plated on selective media (SD media lacking leucine (5 g/L ammonium sulfate, 1.7 g/L yeast nitrogen base without amino acids (MP Biomedicals; 4510522), 1 NaOH pellet (VWR; BDH9292), 0.69 g/L CSM-LEU (amino acid mix lacking leucine), 2% glucose, and 15 g/L Bacto Agar for plates)) that had been filter sterilized prior to seeding. All media except YPD (autoclaved) was 0.2μm filter sterilized.

Phosphate Buffered Saline (PBS) was prepared by creating two individual solutions of mono-and dibasic potassium phosphate (100 mM potassium chloride, 25 mM potassium phosphate). Solutions of different pH values of PBS were prepared individually by mixing the two individual solutions of mono– and dibasic solutions together until the desired pH level was achieved as measured with an Accumet XL150 pH meter (Fisher Scientific).

### Plasmids

The yeast plasmid pYEplac181-pHluorin was provided by the Rajini Rao Lab at Johns Hopkins University. The pHluorin gene from pYEplac181-pHluorin was cloned into the pLIC-His vector (pLIC-His-pHluorin) so that the pHluorin could be expressed recombinantly and purified from E. coli. The mammalian plasmid pcDNA3.1(+)-pHluorin was generated by GenScript (Piscataway, NJ) through synthesizing and cloning a codon optimized version of the pHluorin gene into the pcDNA3.1(+) backbone. Human OPRM, OPRD, and OPRK cDNA from the PRESTO-TANGO library was sub-cloned into the pYEplac181 vector using the NEBuilder HiFi DNA Assembly Kit (NewEnglandBioSciences, E2621S) to create a yeast 2µ high copy plasmid for expressing human mu-OR, delta-OR, and kappa-OR.

All mutants were created from the receptor-respective pYEplac181 backbones using the strategy called the 5’-phosphate PCR Assembly. Briefly, vectors were amplified using Q5 High-Fidelity DNA Polymerase (NEB) reactions with a 5’-phosphate-modified complementary primer, another primer that contains the desired change and a complementary region (facing opposite directions), and the template plasmid DNA. PCR products were verified by agarose gel electrophoresis, then digested with DpnI (NEB) at 37°C for 1.5 hours, followed by heat inactivation. Ligation was performed overnight at 16°C or two hours at room temperature using T4 DNA Ligase (NEB). Ligase was heat-inactivated, and 2µL of the ligation mix was transformed into competent DH-5α cells.

All twenty isoforms were retrieved from Uniprot (P35372), including computationally mapped potential isoform sequences. Then they were codon optimized for yeast using GenSmart (Genscript) and ordered as gblocks (IDTDNA), which were then inserted into the pYEplac181 backbones using the NEBuilder HiFi DNA Assembly Kit (NEB, E2621S).

All plasmids were verified via Sanger or nanopore sequencing (Eurofins Genomics).

### DCyFIR Profiling

*Solutions.* 1× TE buffer (10 mM Tris, 1 mM EDTA). Lithium acetate (LiOAc) mix (100 mM LiOAc, 10 mM Tris, 1 mM EDTA). PEG mixture (40% (w/v) PEG 3350, 100 mM LiOAc, 10 mM Tris, and 1 mM EDTA).

*Strain Generation.* Yeast cells were initially inoculated into 3 mL of YPD medium and incubated overnight at 30 °C with shaking. The following day, 100 µL of this starter culture was transferred into 5 mL of fresh YPD and incubated for approximately 2.5 hours until the culture reached an optical density (OD) of 0.6–1.2. The cells were harvested, and the resulting pellet was washed sequentially with 5 mL of 1× TE buffer and 5 mL of the LiOAc mix. After the final wash, the yeast pellet was resuspended in 200 µL of LiOAc mix to prepare for transformation.

For each transformation reaction, 50 µL of the chemically prepared yeast cells were combined with 300 ng of plasmid DNA, 20 µL of repair DNA or donor payload, 350 µL of PEG mix, and 5 µL of salmon sperm DNA in a sterile 1.5 mL microcentrifuge tube. The samples were briefly vortexed and incubated at room temperature for 30 minutes. Following this incubation, 24 µL of 100% DMSO was added to the mixture, and the cells were subjected to a heat shock at 42 °C for 15 minutes. Post-heat shock, the cells were pelleted by centrifugation at 10,000 rpm for 3 minutes, gently resuspended in 100 µL of YPD without vortexing, and plated onto leucine-depleted selective media for approximately 3 days of growth at 30 °C.

*Strain Growth.* Prior to performing DCyFIR profiling, each A2AR-variant-Gα strain was streaked out on YPD media plates and incubated at 30°C for two days. One colony was picked from each plate and seeded into one milliliter of SCD media contained in a ninety-six-well deep well plate. Cultures were allowed to grow overnight at thirty degrees Celsius in a static environment. The cells were then diluted down to an OD600 of approximately 0.05 at both pH levels and grown to mid-log phase during the day. The cells were then diluted down again to an OD600 that would allow them to have an OD600 value of one when they were harvested twenty-four hours later. This usually corresponds to a 16 to 18 hour period of growth under non-shaking conditions at 30C. It should be noted that there is a difference in doubling time between pH levels that requires a greater initial concentration of cells at pH seven versus pH five. Experimental Setup, Measurements, Replicates. Following the production strain growth procedure described above, the cells were collected and adjusted to an equal concentration based on an OD600 reading of zero point one. These normalized cultures were divided equally among four wells of a three-eighty-well plate as follows: forty-five microliters of each normalized culture per condition was placed within four wells of a three-eighty-well plate to generate four independent replicate sets while generating a total of sixteen data points for each condition: four data points for each vehicle condition at each pH level and four data points for each agonist condition at each pH level. Each well received forty microliters total after adding four microliters of vehicle or ten times dilution of agonist. The cultures were allowed to continue growing for an additional eighteen hours at pH five and twenty-four hours at pH seven until they reached an OD600 value ranging from three to four. Because the rate of growth is slightly less at pH seven than at pH five due to differences in doubling time it was necessary to continue growing some subsets of the receptor population an additional twenty-four hours beyond the eighteen hours required for cells grown at pH five in order for them to reach an OD600 value range of three to four. Therefore, all DCyFIR profiling results shown in Figs. 2, 4, 5 were obtained from OD600 matched cultures. GPCR signaling activity was determined through measurement of fluorescence emitted from the mTq2 transcriptional reporter through quantitative measurement of the fluorescence of the OD-matched cultures utilizing a ClarioStar microplate reader with the following settings: Bottom Read; Ten Flashes/Well; Ex-F: 430 – 10nm; Dichroic Filter: LP458nm; Em-F:482 – 16nm; Gain =1300.

### pH Titration Experiments

Strains were grown. Each GPCR-Gα strain was taken out of a glycerol stock on YPD plates and incubated at 30°C for 48 hours. A single colony for agonist-treated pH titrations was selected and transferred to 5 ml of SCD media at pH 5.0, 5.5, 6.0, 6.5, and 7.0. The same process was followed to select a single colony that would be added to 5 ml of SCD media at pH 5.0 or 7.0 for constitutively active GPCR-Gα pH titrations based upon the level of activation by low or high pH, respectively. Cells grown in each production run were diluted to an optical density at 600 nm of 0.05 (for pH 5.0, 5.5, and 6.0 growths) or 0.1 (for pH 6.5 and 7.0 growths), using each of the different pH media in a 50 ml conical tube, shaken at 30°C until mid-log phase was reached during the day. On the second day, cells were back-diluted so that they will be at an OD600 value of 1.0 after 4-6 hours, with the use of fresh media at the desired time after diluting overnight-shaken cultures at 30°C, so that they are completely adapted to the pH of the media. Then the cultures were subject to DCyFIR profiling methods as mentioned above.

### pHluorin Calibration

A pHluorin standard curve was calculated using purified ratiometric pHluorin. 200 μL PBS buffer titrated to each pH (137 mM NaCl, 2.7 mM KCl, 10 mM Na₂HPO₄, 1.8 mM KH₂PO₄) supplemented with 20 mM MES (Sigma-Aldrich, #M3671) or 20 mM HEPES (Sigma-Aldrich, #H3375, adjusted to the indicated pH values using HCl or NaOH) were placed into a 96-well microplate (CytoOne; CC7626-7596), and 36 μL aliquots were moved in technical quadruplicate to a black bottom 384-well plate (Greiner, 781096). Briefly, 4 μL ratiometric pHluorin (1 μM) were resuspended in each pH PBS buffer. The plate was vortexed at 2000 rpm for 30 seconds before reading. Excitation spectra were collected using a ClarioStar microplate reader (BMG LabTech) with the following parameters: top read, 40 flashes/ well, excitation start: 350 to 10 nm, excitation end: 495 to 10 nm, 5 nm steps; emission: 520 to 10 nm; instrument gain: 1700; focal height 7 mm. Raw fluorescence values at 385 and 475 nm were used to calculate the pHluorin ratio (385/475 nm) for each technical replicate. A standard curve was built by plotting the pHluorin ratio as a function of pH and fit using a sigmoidal 4PL model (X = log[concentration]) in GraphPad Prism. Experimental error was calculated in GraphPad Prism as the SD of the mean for the n=4 technical replicates.

### Acid Stress Assay and Phenotypic Classification

For acid stress assays, the HEK293T cells was seeded into the aforementioned white Greiner 384-well plates at a density of 5,000–10,000 cells per well and allowed to adhere for 48 h post-transfection or post-transduction. Cells were washed with pre-warmed PBS and incubated in pH-adjusted buffer (137 mM NaCl, 2.7 mM KCl, 10 mM Na₂HPO₄, 1.8 mM KH₂PO₄) supplemented with 20 mM MES (Sigma-Aldrich, #M3671) or 20 mM HEPES (Sigma-Aldrich, #H3375, adjusted to the indicated pH values using HCl or NaOH)) for 10 minutes at room temperature. pH values ranged from 5.0 to 7.0 in 0.5-unit increments. Each well received 30 µL of buffer at the indicated pH.

Excitation spectra were collected using a ClarioStar microplate reader (BMG LabTech) with the following parameters: top read, 40 flashes/ well, excitation start: 385 to 10 nm, excitation end: 475 to 10 nm, 15 nm steps; emission: 520 to 10 nm; instrument gain: 2750; focal height 7 mm. Raw fluorescence values at 385 and 475 nm were used to calculate the pHluorin ratio (385/475 nm) for each technical replicate.

## Author contributions

D.G.I. managed the study, and D.G.I. and K.D.L. wrote the manuscript. K.D.L., A.W., G.S., B.G., and S.T. performed all experiments.

## Acknowledgements

This work was supported by the National Institutes of Health through an R35 Maximizing Investigators’ Research Award (R35GM119518 to D.G.I.), which provided core funding for this study. Additional support was provided by a Pap Corps Champions for Cancer Research Endowed Chair to D.G.I. and the Sylvester Comprehensive Cancer Center.

